# Uneven terrain versus dual-task walking: differential challenges imposed on walking behavior in older adults are predicted by cognitive and sensorimotor function

**DOI:** 10.1101/2023.03.14.531779

**Authors:** Valay A Shah, Yenisel Cruz-Almeida, Arkaprava Roy, Erta Cenko, Ryan J Downey, Daniel P Ferris, Chris J Hass, Patricia A. Reuter-Lorenz, David J Clark, Todd M Manini, Rachael D Seidler

## Abstract

Aging is associated with declines in walking function. To understand these mobility declines, many studies have obtained measurements while participants walk on flat surfaces in laboratory settings during concurrent cognitive task performance (dual-tasking). This may not adequately capture the real-world challenges of walking at home and around the community. Here, we hypothesized that uneven terrains in the walking path impose differential changes to walking speed compared to dual-task walking. We also hypothesized that changes in walking speed resulting from uneven terrains will be better predicted by sensorimotor function than cognitive function. Sixty-three community-dwelling older adults (65-93 yrs old) performed overground walking under varying walking conditions. Older adults were classified into two mobility function groups based on scores of the Short Physical Performance Battery. They performed uneven terrain walking across four surface conditions (Flat, Low, Medium, and High unevenness) and performed single and verbal dual-task walking on flat ground. Participants also underwent a battery of cognitive (cognitive flexibility, working memory, inhibition) and sensorimotor testing (grip strength, 2-pt discrimination, pressure pain threshold). Our results showed that walking speed decreased during both dual-task walking and across uneven terrain walking conditions compared to walking on flat terrain. Participants with lower mobility function had even greater decreases in uneven terrain walking speeds. The change in uneven terrain speed was associated with attention and inhibitory function. Changes in both dual-task and uneven terrain walking speeds were associated with 2-point tactile discrimination. This study further documents associations between mobility, executive functions, and somatosensation, highlights the differential costs to walking imposed by uneven terrains, and identifies that older adults with lower mobility function are more likely to experience these changes to walking function.

## 1 Introduction

Even “healthy” aging often leads to deficits in cognitive, sensory, and motor processes. These changes have been associated with age-related declines in walking function (Paraskevoudi et al., 2018). Cognitive changes can impact the top-down control of walking (for instance, attentional modulation of sensory inputs), while sensory changes are thought to impact the bottom-up, feedback-based control of walking. Walking ability is also considered to be an indicator of overall health and poor walking function (e.g., decreased speed; increased gait variability) is associated with higher morbidity and fall risk (Manini et al., 2017; Wilson et al., 2019). These mobility declines decrease quality of life and independence, and increase overall healthcare costs (Hardy et al., 2011; Knaggs et al., 2011; Newman et al., 2006). Thus, understanding and quantifying walking behavior as we age is essential for developing interventions and reducing the burden of walking deficits.

With age, walking becomes less automatic and more dependent on cognitive control (Clark, 2015; Fettrow et al., 2021; Hausdorff et al., 2005; Yogev-Seligmann et al., 2008). Higher-order cognitive processes such as attention and executive function (Reuter-Lorenz et al., 2016) that activate prefrontal brain regions (Holtzer et al., 2011; Pizzamiglio et al., 2017) are engaged to enhance/maintain walking performance. Studies have shown that older adults with compromised executive and attentional function have poorer walking capability (Atkinson et al., 2007; Ble et al., 2005; Callisaya et al., 2015; Holtzer et al., 2006; Watson et al., 2010). Increased involvement of cognitive processes in older adults is also evident by robust decrements in walking behavior, such as slower speed and greater variability in stride length, compared to younger adults when walking is performed concurrently with a cognitive task [e.g., word generation or verbal fluency (Beurskens et al., 2014; Blumen et al., 2014; IJmker & Lamoth, 2012); arithmetic (Al-Yahya et al., 2016; Springer et al., 2006); word recall (Lindenberger et al., 2000); visual cue detection (Beurskens & Bock, 2013; Protzak et al., 2021)]. These dual-task experiments have provided valuable insights into the role of cognition in walking and have contributed greatly to our understanding of age differences in responses to increasing walking task difficulty. However, dual-task walking paradigms alone do not fully address the challenges faced by older adults in the community and environments where fall risk is increased.

Community-dwelling older adults have a greater risk of falls when walking around the home and community, where uneven terrains are frequently encountered [e.g., walking over a rug, walking from grass to concrete (Ambrose et al., 2013; Berg et al., 1997; Kelsey et al., 2012)]. Compared to dual-task paradigms, an approach that examines walking ability through uneven terrains may better mimic walking environments where fall incidents are prevalent, and therefore have greater ecological relevance. Previous studies have manipulated walking task difficulty by changing path width, adding obstacles, or with uneven terrains/surfaces in the walking path (Chatterjee et al., 2020; Darici & Kuo, 2022; Downey et al., 2022; Kotegawa et al., 2021; Marigold & Patla, 2008a, 2008b; Voloshina et al., 2013). Marigold and Patla (2008a) investigated the role of vision and age differences in gait speed and step length and width during multi-surface terrain walking. Their results showed that older adults had decreased step length relative to young adults while also having slower gait speed when walking across multi-surface terrains compared to flat surface walking. Additionally, older adults showed an increased reliance on vision and increased head pitch when vision from the lower visual field was blocked across flat and multi-surface terrains. In a recent study, Kotegawa et al. (2021) increased walking difficulty by narrowing the width of the walking path in older adults. This increased walking difficulty resulted in greater time needed to navigate the path (i.e., decreased walking performance). Given the ubiquity of uneven surfaces in everyday environments (Chen et al., 2015; Patla & Shumway-Cook, 1999; Rantakokko et al., 2013) and the ecological relevance of uneven terrains for fall risk, we sought to extend these previous findings by parametrically manipulating the extent of terrain unevenness in the walking path. Parametrically varying the unevenness of the pedestrian terrain has the potential to also delineate the sensorimotor versus cognitive control of walking. Varying the difficulty of the walking task may challenge both the vestibular and proprioceptive systems differently from any challenges caused by age-related changes to cognition.

In the present study, we quantified the cost (i.e., change in walking behavior) associated with dual-task walking in comparison to walking on varying levels of uneven terrains in older adults. We analyzed data collected from participants in the Mind in Motion study (Clark et al., 2020). Older adult participants performed uneven terrain and dual-task overground walking. We quantified changes in walking speed as walking difficulty increased by either uneven terrains or a dual-task. We hypothesized that: 1) uneven terrains in the walking path will impose differential changes to walking speed compared with dual-task walking, and 2) changes in walking speed resulting from uneven terrains will be better predicted by sensorimotor function (e.g., pain thresholds) than cognitive function. Characterizing the effects and predictors of different types of walking challenges for older adults can advance understanding of the causes and consequences of walking declines in this population. Such characterization can also identify subpopulations of older adults that would benefit from future rehabilitation and training interventions (cognitive or physical) to reduce falls.

## 2 Methods

We analyzed data from 63 community dwelling, older adults (76 ± 6.59 yrs [mean ± SEM here and onwards], 32 males) who participated in the Mind in Motion study. Older adults were based in the north central Florida community and provided written informed consent to complete experimental procedures approved by the University of Florida Institutional Review Board. General inclusion criteria included: >=65 years of age, capability to walk 400m within 15 minutes without sitting, no brain injuries (stroke, concussion), no major hospitalizations in previous 6 months, no use of a walker or wheelchair, and eligibility for Magnetic Resonance Imaging (for a detailed version of the inclusion and exclusion criteria see Supplemental Table 1). The cohort of older adults included a wide range of mobility function, assessed via the Short Physical Performance Battery test [SPPB (Guralnik et al., 1994)], making our sample more representative of older adults in the community. The SPPB combines measurements of the ability to stand for up to 10 seconds with feet positioned in three ways (together side-by-side, semi-tandem and tandem); time to complete a 3-m or 4-m walk; and time to rise from a chair. This gives a comprehensive score of mobility function (ranging from 0-16, with 16 being highly mobile). Older adults with SPPB >= 10 were placed in a high functioning (HFOA) mobility group and those with SPPB <10 were placed in a low functioning (LFOA) mobility group.

### 2.1 Participant Visit

During a study visit, participants completed sensory, motor, and cognitive assessments. We collected sensory data using the two-point discrimination test and pressure pain threshold, and sensorimotor data via grip strength and walking tests. Cognitive assessments were performed using a subset of tests from the NIH toolbox for Cognition. Within the same study visit, participants also performed overground uneven terrain walking and verbal dual-task walking as described below.

#### 2.1.a Uneven Terrain Walking

Participants performed overground walking on uneven terrain mats. Uneven terrain walking was performed with four terrain unevenness levels (Flat, Low, Medium, and High terrain unevenness). Terrain unevenness was modified via rigid foam disks that were attached on a 3.5 m walking mat placed on the ground. The Flat condition had green circles painted on the mat (i.e., normal walking). The Low condition consisted of yellow-colored 1.3 cm-high disks. The Medium (Med) condition had orange-colored disks of two heights: 50% were 1.3 cm and 50% were 2.5 cm. The High conditions had red-colored disks of three heights: 20% were 1.3 cm, 30% were 2.5 cm, and 50% were 3.8 cm. Figure 1 shows the walking mats with varying uneven terrain levels. People were informed about the unevenness of the walking path and the corresponding colors. Participants were instructed to walk at a normal, comfortable pace and wore their own shoes. They walked on each level of uneven terrain three times and the time to walk the middle 3-meter portion of the walking mat was measured via a stopwatch.

**Figure 1:**
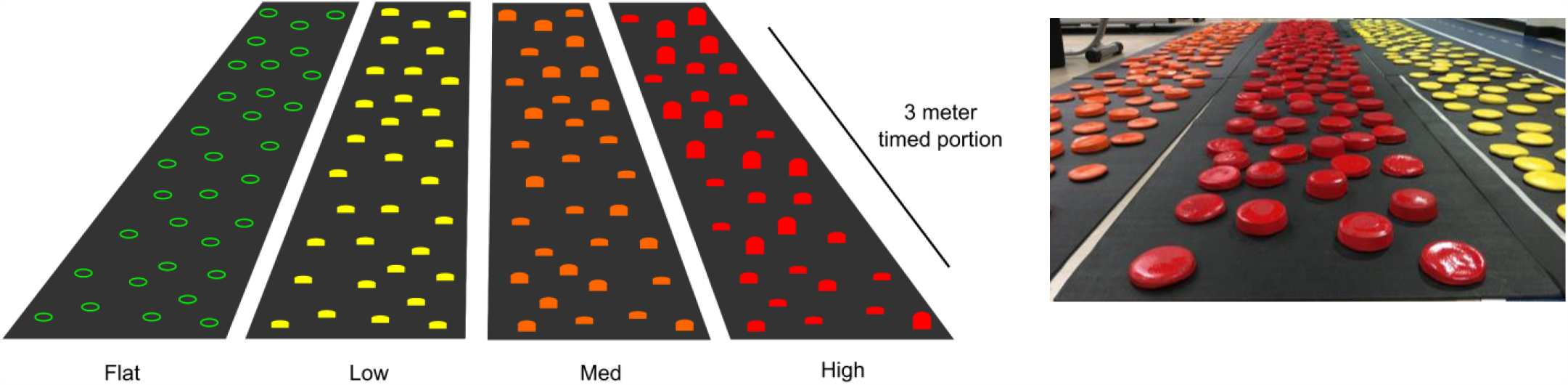
Schematic and photo of uneven terrain walking mats. Green circles: Flat terrain (no uneven terrain), Yellow disks: Low condition (1.3 cm-high disks). Orange disks: Medium condition (1.3 cm and 2.5 cm disks). Red disks: High condition (1.3 cm, 2.5 cm, and 3.8 cm disks).

#### 2.1.b Verbal Dual-task Walking

Older adults completed single- and dual-task walking on a 4.6 m GAITRite walkway with embedded pressure-activated sensors (CIR Systems). For the single-task, participants walked on the GAITRite mat at a normal, comfortable pace, while wearing shoes. Participants were instructed to repeat this task three times. For the dual-task, participants were instructed to walk at a normal, comfortable pace with shoes while concurrently stating as many words as possible starting with a selected letter (e.g., verbalize words starting with the letter R, while walking). The word generation letter was selected by a study coordinator from one of the following seven letters: E,G,I,L,N,O,R. The study coordinator announced the letter concurrently with the first step of the participant in the GAITRite mat. Responses were recorded to calculate verbal fluency and dual task costs. Duplicate words were not counted towards verbal fluency (e.g., “Red, Read, Road, Red, Rough” resulted in 4 words generated). Three walking trials were completed for the dual-task walking, with each trial having a new letter selected for word generation.

#### 2.1.c Sensorimotor and Cognitive Testing

Two-point discrimination (2PT Disc) was tested on the non-dominant foot on the plantar surface of the head of the first metatarsal. Foot dominance was self-reported by the participant. A calibrated two-point discrimination device was used to measure two-point thresholds with 15, 13, 7, and 4 mm two-point distances. The participant laid supine on a medical examination table and their view of the foot was occluded. Starting from the largest distance, we tested each two-point distance for five trials: twice with the points oriented vertically, twice horizontally, and once with just a single point (the trials were conducted in a random order for each distance). Participants were instructed to verbally indicate when they felt the device touch the skin and whether it was with one or two points. No other instructions were given. For each correct indication participants received a score of 1, and for false-positive or false negative indications they received a score of 0. We continued the test to the next smallest separation only if the score was 3 or larger. The threshold was determined as the smallest distance where the participant correctly identified the number of contact points in at least three of the five trials.

Pressure pain threshold (PPT) was measured on the right thigh (midway between the knee and hip) over the quadriceps muscle via an algometer (Wagner Instruments: model FDX 25). Participants were seated in a relaxed, upright position in a chair, with both arms supported by the arm rests. The tip of the algometer was placed perpendicular to the skin over the thigh and pressed down. Participants were instructed to verbally indicate when the algometer tip became painful and the threshold was recorded in kPa. Participants completed three to five trials of the pressure pain measures. We aimed to record three PPT values, however, when the measurements differed by more than 40 kPa, we proceeded with the completion of five trials. The PPT was determined as the average of all trials.

Grip strength was measured for the dominant hand via a handgrip dynamometer (Jamar: Model 63785). We used grip strength as a measure of overall motor performance because it is highly associated with mobility function (Bohannon, 2019; Carson, 2018; Pratama & Setiati, 2018; Rantanen et al., 1999). Participants gripped the dynamometer while sitting in a chair with their arm resting on a table, with 90° elbow flexion. If participants indicated worsened arthritic pain in the dominant hand or had undergone hand or wrist surgery, we measured grip strength for the non-dominant hand. Participants were instructed to grip the dynamometer and squeeze as hard as possible. After a practice trial to get familiarized with the device, participants performed two trials and the maximum force (N) was recorded from the dynamometer. Grip strength was determined as the average of the two trials.

We assessed cognitive function using a subset of tests from the NIH toolbox cognitive battery (Heaton et al., 2014; Weintraub et al., 2013). Participants completed the Dimensional Change Card Sort test (DCCS), The List Sorting Working Memory test (LS), and a Flanker test. The Dimensional Change Card Sort task provided a measure of cognitive flexibility (Zelazo, 2006; Zelazo et al., 2014). Participants first matched test images to one feature of target images (e.g., color). Occasional switch trials asked participants to match test images to another feature of the target images (e.g., shape). The List Sorting Working Memory test provided a measure of working memory capacity and recall (Tulsky et al., 2013). Participants were presented with a sequence of stimuli (i.e. illustrations of an animal, object, food item, etc.), presented both visually and aurally on an Ipad (Apple Inc). A picture of each stimulus was displayed for two seconds while the name of the stimulus was simultaneously read via a computerized voice. Participants were required to remember each stimulus, mentally reorder them from smallest to largest in size, and verbalize the stimuli in this order. The Flanker task provided a measure of attention and inhibitory function (Eriksen & Eriksen, 1974; Zelazo et al., 2014). Participants were required to indicate the left or right pointing direction of a central stimulus while ignoring the incongruent stimuli around it (i.e., the flankers, typically two on either side). We used the computed score (combination for task accuracy and reaction time) outputs for the NIH DCCS and Flanker task (range 0-10), and the raw score (no computed score is calculated for task accuracy) for the NIH LS test (range 0-26) output by the NIH toolbox software. We chose to use the computed and raw scores rather than the age-corrected standard scores as we wanted to gauge the absolute performance in each test for each participant, rather than comparing them to the NIH Toolbox nationally representative normative sample. The interpretation of the computed and raw scores as absolute performance follows the guidelines of the NIH Toolbox Scoring and Interpretation guide.

### 2.2 Statistical Analyses

To better understand the data distribution in our cognitive and sensorimotor predictors, we performed correlations across our predictors with the two older adult groups combined and independent samples t-test on the predictors to determine whether they differed between HFOA and LFOA. During dual-task walking, verbal fluency was calculated as the average of the number of words generated across three verbal dual-task walking trials and compared across HFOA and LFOA using independent samples t-test. We also correlated verbal fluency with percent change in walking speed between single and dual-task walking to determine dual-task related costs to walking behavior. We then tested our hypotheses that uneven terrains in the walking path will impose differential changes to walking speed than dual-task walking and that changes in walking speed resulting from uneven terrains will be better predicted by measures of sensorimotor function than cognitive function. We first calculated percent change in walking speed from the flat, overground walking condition (Flat vs. Low, Flat vs. Med, Flat vs. High unevenness, and single vs. dual-task). We will refer to these percent speed changes as Flat-Low, Flat-Med, Flat-High, and Single-Dual. To compare walking performance across tasks and groups, we performed a mixed-model ANOVA, with task level (Low, Med, High terrain unevenness, and dual-tasking) as the within-subject fixed factor and mobility function group (HFOA and LFOA) as a between-subjects fixed factor. Percent change in walking speed was the dependent variable. We used post-hoc Holm-Bonferroni corrected t-tests to compare changes in walking speed across tasks and groups. We then examined whether sensorimotor and cognitive function scores predicted percent changes across uneven terrain and dual-task walking in a similar manner. We used separate linear regression models to investigate sensorimotor (2PT-Disc, PPT, and grip strength) and cognitive (NIH LS, NIH DCCS, and NIH Flanker) predictors of percent change in walking speed. We separated regression models for uneven terrain walking (Flat vs. High unevenness) and dual-task walking (single vs. dual-task). We then included SPPB scores and sex as covariates in the models to account for mobility function differences and sex differences. We examined the variance inflation factor to assess multicollinearity in the regression models. Statistical significance was set at a family-wise error rate of α = 0.05.

## 3 Results

### 3.1 Participants and Behavioral Performance

Our sample cohort of older adults included 26 high functioning older adults (HFOA; 74.6 ± 3.7 yrs, 20 males; SPPB = 11 ± 0.94) and 37 low functioning older adults (LFOA; 76.9 ± 7.8 yrs, 12 males; SPPB = 7.77 ± 1.27). As expected, mobility function was different between the two older adult groups, which were prospectively separated based on their SPPB scores (t = 11.2, p<0.001). There were no significant differences in sensorimotor scores between these mobility function groups, but NIH LS and DCCS scores were lower for LFOA compared to HFOA (p<0.05). Across all older adults in our study, we found significant correlations within the cognitive scores, where NIH DCCS scores were correlated with NIH LS (r = 0.494, p<0.001) and NIH Flanker (r = 0.648, p<0.001) scores. Sensorimotor scores were also correlated in that PPT was correlated with grip strength (r = 0.394, p=0.002) and 2PT-disc (r = 0.227, p=0.039) scores. Table 1 shows the average sensorimotor and cognitive scores and Table 2 shows the groupwise scores for our measurements. Although we did not perform specific statistical tests for differences across task conditions, we also noted a decrease in walking speed as walking complexity increased both with uneven terrains in the walking path and dual-task walking. Figure 2 shows the distribution of walking speeds across all older adults. Interestingly, the Flat walking speed (0.903 ± 0.037) was lower than single-task walking speed (1.11 ± 0.028). Thus, the percent change in uneven terrain and dual-task walking speed from the Flat and single-task walking speeds are justified rather comparisons of absolute speed changes across the two tasks. During dual-tasking, HFOA generated on average 4.60 ± 0.256 words while LFOA generated 3.93 ± 0.192 words while walking and the difference between groups in their verbal fluency during walking was significant (t = 2.14, p=0.036). Verbal fluency was also correlated with NIH LS task (r = 0.314, p=0.019) as well as Single-Dual percent speed change (r = -0.361, p=0.004). The correlation between percent speed change and verbal fluency was stronger for LFOA (r = -0.560, p<0.001) compared to HFOA (r = -0.240, p=0.248). The negative correlation suggests that older adults, especially LFAO, who generate more words during dual-task walking, do so at a cost to walking speed.

**Table 1:** Sensorimotor scores and cognitive scores (mean and SEM) for older adult participants. LS = List Sorting Working Memory; DCCS = Dimension Change Card Sort; PPT = Pressure Pain Threshold; Significantly correlated with each other: *,τ,**,ττ = p<0.05.

| Measure | Score |
| --- | --- |
| NIH LS | 16.1 (0.3)* |
| NIH DCCS | 7.46 (0.12)*,$\tau$ |
| NIH Flanker | 7.46 (0.10)$\tau$ |
| PPT (kPa) | 37.3 (3.1)**,$\tau\tau$ |
| Grip Strength (N) | 23.6 (1.6)** |
| 2PT Disc (mm) | 12.4 (0.4)$\tau\tau$ |

**Table 2:**
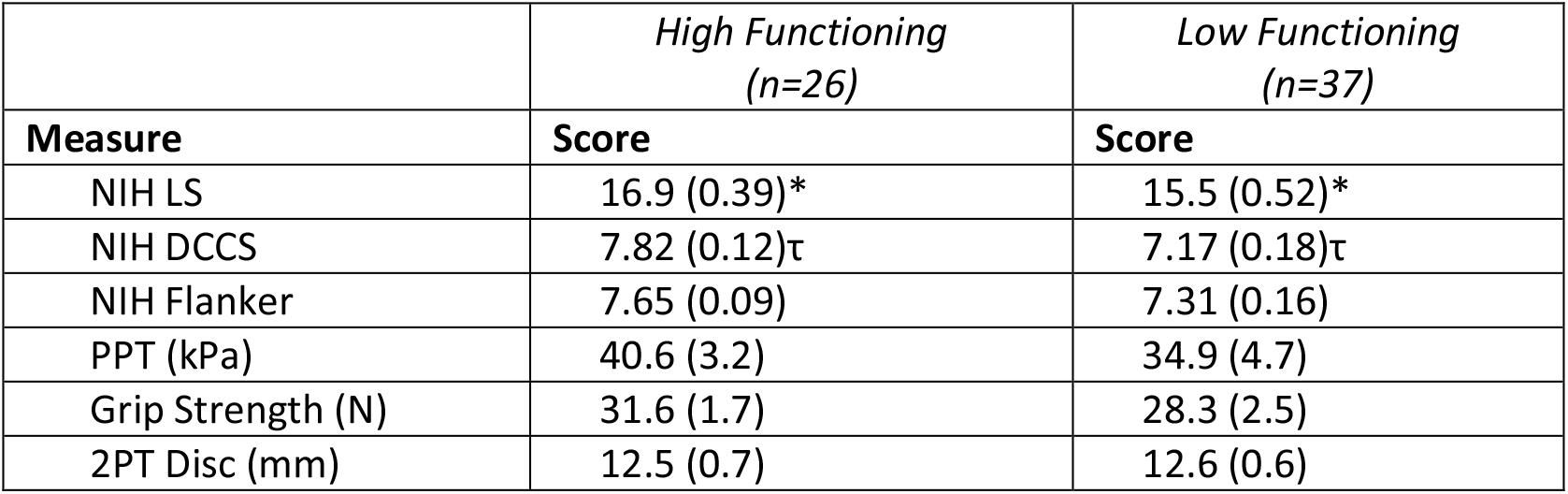
Sensorimotor and cognitive scores (mean and SEM) separated by SPPB based mobility function. ; Significantly different: *,τ = p<0.05.

|  | *High Functioning*<br>*(n=26)* | *Low Functioning*<br>*(n=37)* |
| --- | --- | --- |
| Measure | Score | Score |
| NIH LS | 16.9 (0.39)* | 15.5 (0.52)* |
| NIH DCCS | 7.82 (0.12)$\tau$ | 7.17 (0.18)$\tau$ |
| NIH Flanker | 7.65 (0.09) | 7.31 (0.16) |
| PPT (kPa) | 40.6 (3.2) | 34.9 (4.7) |
| Grip Strength (N) | 31.6 (1.7) | 28.3 (2.5) |
| 2PT Disc (mm) | 12.5 (0.7) | 12.6 (0.6) |

**Figure 2:**
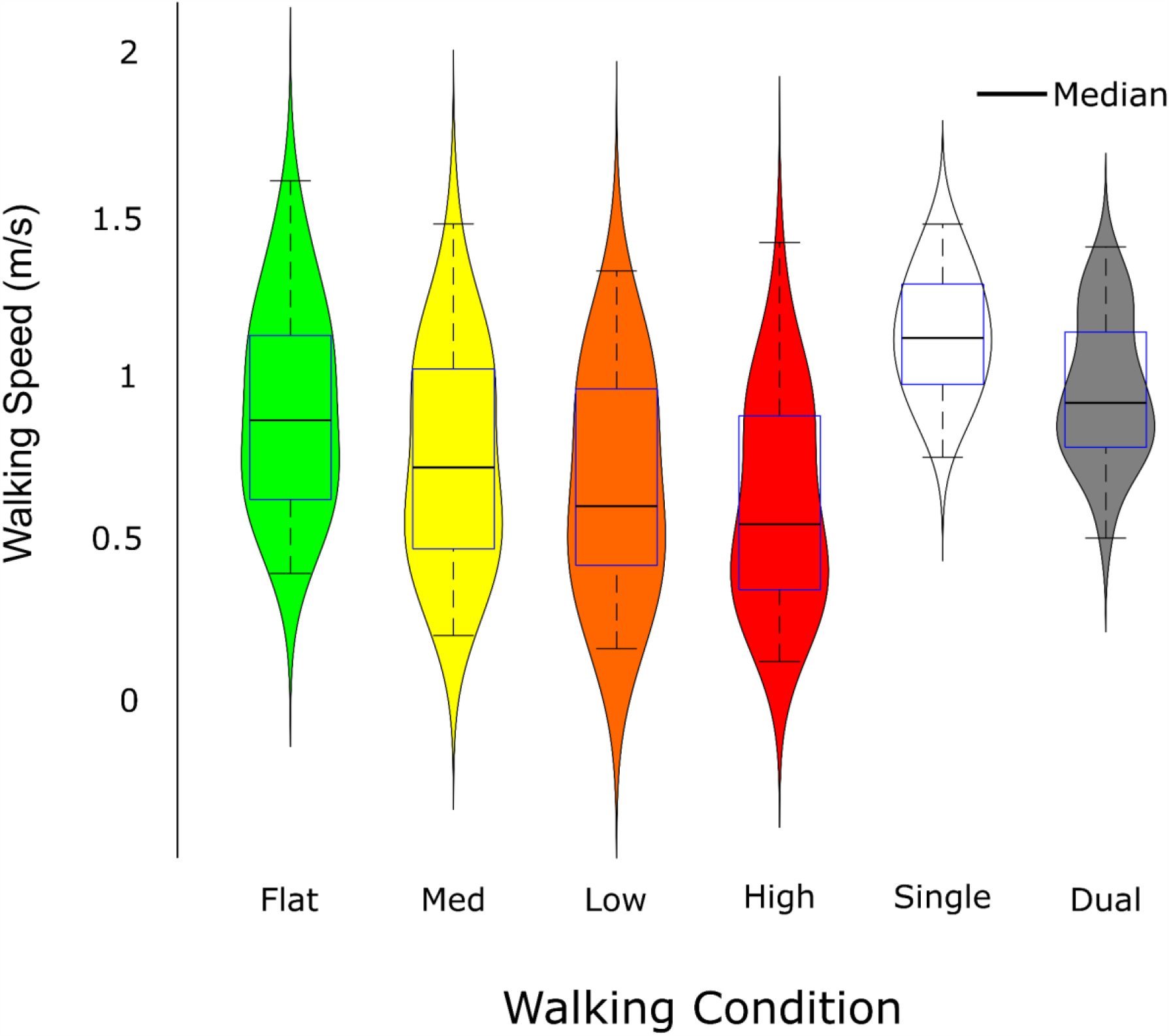
Violin plots showing the distribution of the self-selected overground walking speeds during uneven terrain and dual-task walking conditions, across all older adults. Statistical tests were not performed but as walking complexity increased, either via uneven terrains or with a verbal dual-task, walking speed decreased.

### 3.2 Percent Changes in Walking Speed

We performed a check on data normality with a Shapiro-Wilk test on percent walking speed change (Flat-High and Single-Dual conditions) which resulted in p>0.05, indicating that the distribution of the data was not significantly different from the normal distribution. Figure 3 shows the change in speed across the two groups and task conditions. A mixed-model ANOVA on percent change in walking speed resulted in a significant effect of task level (F_(3,183)_ = 39.2, p<0.001, η^2^ =0.39; Fig 3: Horizontal Blue Lines). Post hoc multiple comparisons (Holm-Bonferroni corrected) revealed significant differences in the percent speed change across tasks (Single-Dual vs. Flat-Med, p<0.001, Cohen’s d=0.70; Single-Dual vs. Flat-High, p<0.001, d=1.0; Flat-Low vs. Flat-Med, p<0.001, d=0.78; Flat-Low vs. Flat-High, p<0.001, d=1.3; Flat-Med vs. Flat-High, p<0.001, d=0.68). As expected, mobility function group also had a significant effect on percent speed change (F_(1,61)_ = 18.1, p<0.001, η^2^ =0.23). Across task conditions, LFOA had more decreases in walking speed (-30.2 ± 1.95 %) compared to HFOA (-17.2 ± 2.32 %). There was also a significant interaction effect between task level and mobility function (F_(3,183)_ = 5.34, p=.002, η^2^ =0.08). This task by mobility function interaction was reflected by LFOA having greater decreases in walking speeds with increasing uneven terrain levels than HFOA. Single-Dual speed change did not differ by mobility function (p=0.187). However, the groups responded differently to uneven terrain in that the LFOA group showed more slowing with each increment of difficulty compared to the HFOA (Flat-Low, p<0.001, d=0.90; Flat-Med, p<0.001, d=0.97; Flat-High, p<0.001, d=1.1; Fig 3: *, **, *** markers).

**Figure 3:**
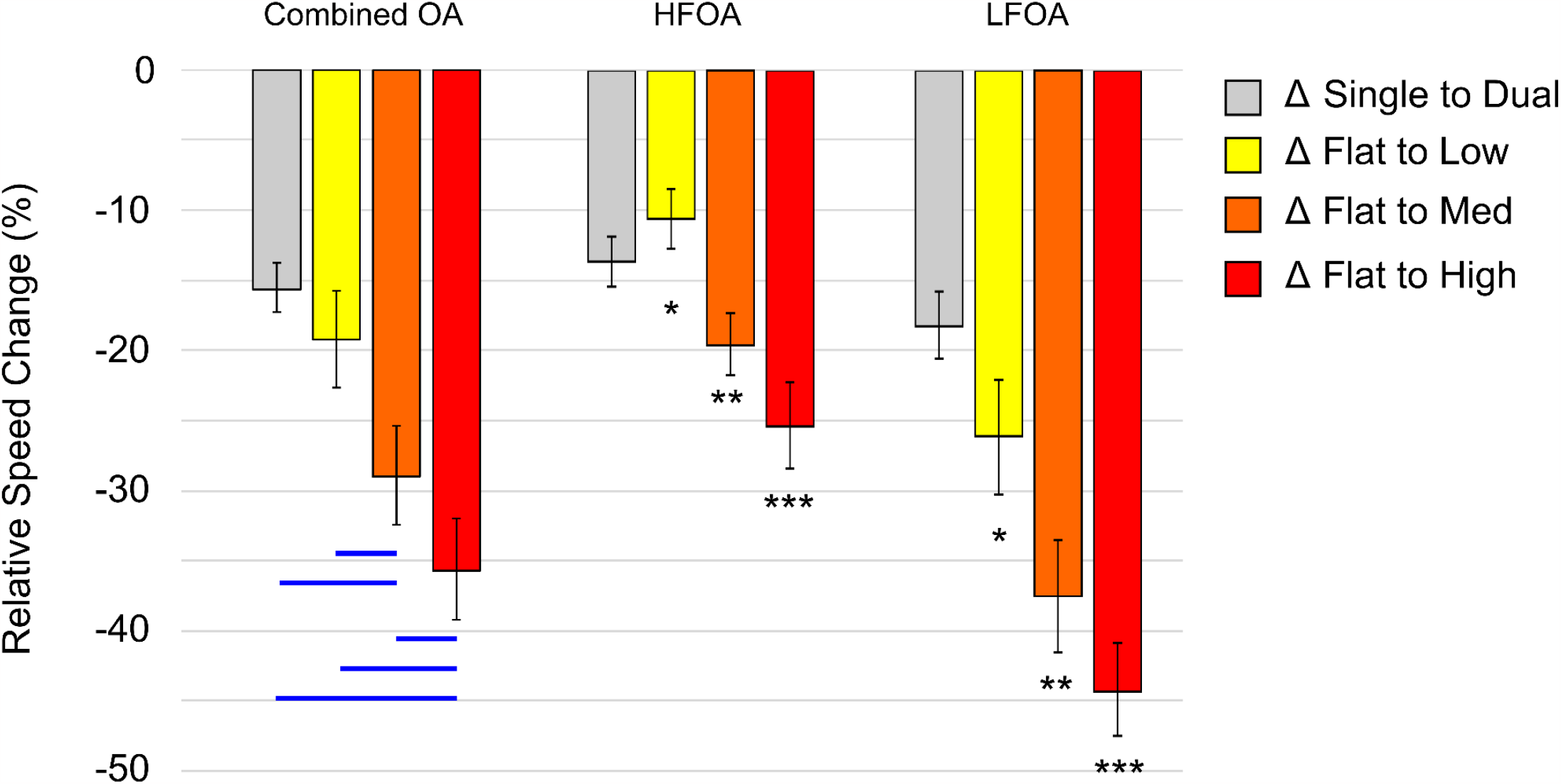
Percent walking speed change (mean and SEM) in older adults across dual-task walking and uneven terrain walking. Gray Bars: single to dual-task speed change. Yellow Bars: Flat to Low terrain speed change. Orange: Flat to Medium terrain speed change. Red: Flat to High terrain speed change. Horizontal Blue lines indicate within group differences (p<0.05). *, **, *** indicate between group differences (p<0.05).

### 3.3 Predictors of Percent Change in Walking Speed

We performed separate multiple linear regression models with percent change in walking speed (Flat-High and Single-Dual walking speed change) as the outcome variable and sensorimotor (grip strength, PPT, 2PT Disc) and cognitive (NIH: DCCS, LS, Flanker) scores as the predictors. No multicollinearity was detected as evidenced by variance inflation factor values for our predictors (NIH DCCS = 2.67, NIH LS = 1.46, NIH Flanker = 1.89, Grip Strength = 1.43, 2PT Disc = 1.26, PPT = 1.26). Six participants were excluded from the cognitive models and seven participants from sensorimotor models due to missing data for one or more predictors. The cognitive regression models were statistically significant for both Flat-High speed change (R^2^ = 0.751, F_(3,53)_ = 53.2, p<0.001) and Single-Dual speed change (R^2^ = 0.704, F_(3,53)_ = 42.0, p<0.001). The NIH Flanker score was a significant predictor of Flat-High speed change (β = -7.76; p=0.049), indicating that lower NIH Flanker scores were related to larger decreases in walking speed across the uneven terrain walking task. The sensorimotor regression models were also statistically significant for both Flat-High speed change (R^2^ = 0.723, F_(3,54)_ = 47.0, p<0.001) and Single-Dual speed change (R^2^ = 0.700, F_(3,54)_ = 42.2, p<0.001). The models found 2PT Disc as a significant predictor for both Flat-High (β = -2.15; p<0.001) and Single-Dual (β = -1.02; p<0.001) speed change, indicating that worse 2PT discrimination function was related to larger decreases in walking speed.

We then added SPPB scores as a covariate to determine the influence of mobility function on predictors for percent changes in speed. The cognitive regression models remained significant for both Flat-High speed change (R^2^ = 0.782, F_4,52)_ = 46.7, p<0.001) and Single-Dual speed change (R^2^ = 0.706, F_(4,52)_ = 31.3, p<0.001). NIH Flanker remained a significant predictor (β = -9.49; p=0.013) for Flat-High speed change even when mobility function was accounted for with SPPB scores. The sensorimotor regression models also remained significant with SPPB as a covariate, for both Flat-High speed change (R^2^ = 0.724, F_4,53)_ = 34.7, p<0.001) and Single-Dual speed change (R^2^ = 0.704, F_(4,53)_ = 31.6, p<0.001). 2PT Disc again remained a significant predictor of both Flat-High (β = -1.94; p=0.007) and Single-Dual (β = -0.837; p=0.023) speed change when mobility function was considered.

We also explored the potential influence of sex on cognitive and sensorimotor predictors. We added sex as a covariate to the cognitive and sensorimotor models, in addition to the SPPB scores. The cognitive regression models were again significant for both Flat-High speed change (R^2^ = 0.793, F_5,51)_ = 39.1, p<0.001) and Single-Dual speed change (R^2^ = 0.709, F_(5,51)_ = 31.3, p<0.001). NIH Flanker score still remained a significant predictor for only Flat-High speed change (β = -8.36; p=0.028). The sensorimotor regression models also remained significant with sex as a covariate, for both Flat-High speed change (R^2^ = 0.745, F_5,52)_ = 30.4, p<0.001) and Single-Dual speed change (R^2^ = 0.756, F_(5,52)_ = 32.3, p<0.001). 2PT Disc again remained a significant predictor of both Flat-High (β = -1.82; p=0.010) and Single-Dual (β = -0.747; p=0.027) speed change. Full results of the regression tables are reported in the Supplemental materials.

## 4 Discussion

The purpose of the current study was to determine and compare performance changes in self-paced walking due to uneven terrains and dual-tasking and how such changes are related to sensorimotor (our measures of 2PT Disc, PPT, and grip strength) and cognitive function (our measures of attention, working memory, and inhibition). We hypothesized that: 1) uneven terrains in the walking path will impose differential changes to walking speed compared with dual-task walking, and 2) changes in walking speed resulting from uneven terrains will be better predicted by sensorimotor function than cognitive function. Our results demonstrated that uneven terrains indeed imposed different costs (i.e., changes) to walking speed, especially when terrain was highly uneven, compared to dual-task walking. Further, whereas dual-task costs to walking speed did not differ between the mobility function groups, costs to walking speed imposed by uneven terrains differentially affected older adults based on their mobility function. Walking speed decreased significantly across both high (HFOA) and low (LFOA) mobility function groups as terrain unevenness increased, however LFOA showed almost a two-fold decrease in walking speed compared to HFOA (Fig 3: HFOA vs. LFOA red bars, indicated by ***). Even the Low level of terrain unevenness imposed almost a two-fold decrease in walking speed in LFOA (Fig 3: HFOA vs LFOA yellow bars, indicated by *). In contrast to uneven terrain walking, Single-Dual walking speed changes did not statistically differ between HFOA and LFOA, although LFOA exhibited greater slowing relative to baseline (Fig 3: HFOA vs LFOA gray bars). Performance of the concurrent word generation (verbal fluency) task, however, differed significantly between HFOA and LFAO, with HFOA generating more words on average while walking. Better word generation by the HFOA group in the absence of greater dual-task cost to walking speed relative to the LFOA suggests several possible differences between the two groups. First, the HFOA may have better baseline functioning in verbal fluency, a possibility supported by the fact that word generation correlates with NIH-LS and by finding that the HFOA performed better on this standardized task than the LFOA group (See Table 2). Second, the HFOA may generally have more executive resources available to allocate to both tasks; or third, because HFOA mobility is generally superior to the LFOA group, walking requires less attention (fewer resources) for the HFOA compared to the LFOA group, allowing more to be devoted to a secondary verbal task. Although the current data don’t permit adjudication among these possibilities, they are not incompatible, and may each play a role in the word generation performance difference we observed. Nevertheless, both groups showed negative correlations between number of words generated and Single-Dual percent speed decrease, indicating that those who produced more words did so at greater cost during dual-task walking. This inverse relationship between magnitude of slowing and words generated was more pronounced for the LFOA, which aligns with the interpretation that this group generally requires greater attention to walking, which is compounded by the demands of doing two things at once. Overall, these findings indicate differential costs between uneven terrain walking and dual-task walking, especially when mobility function is accounted for, an effect we discuss in more detail subsequently.

Both sensorimotor and cognitive models showed that our functional measures accounted for >=70 % of the variability in the change in uneven terrain walking speed. However, this finding contradicts our hypothesis that changes to uneven terrain walking behavior is better predicted by sensorimotor measures. Our sensorimotor regression models showed that 2PT discrimination significantly predicted changes in both uneven terrain and dual-task walking speeds. Older adults who had better 2PT discrimination thresholds also had less changes in walking speeds when walking difficulty was increased. This association between 2PT discrimination and change in walking speed was still significant when we accounted for mobility function and sex differences. These results are in line with previous findings from various studies relating sensory function to walking behavior and partly corroborate our hypothesis that changes to uneven terrain walking behavior is predicted by sensorimotor function. Cutaneous and proprioceptive feedback from the foot and ankle play an important role in the control of walking (Cruz-Almeida et al., 2014; Höhne et al., 2012; Perry et al., 2000). Tactile feedback is also directly used by spinal interneurons and the brain to modulate walking function (Frigon & Rossignol, 2006; Höhne et al., 2012; Hultborn & Nielsen, 2007). Age-related degradations of the peripheral sensory systems also impact the fidelity of sensory feedback used for the cognitive control of walking (Jones & Noppeney, 2021). As a result, older adults may reduce walking speed to improve locomotion coordination and provide more time for the processing of the degraded somatosensory and visual feedback (Coppin et al., 2006; Patla & Vickers, 2003). Interestingly, we did not find any significant association of grip strength or pressure pain threshold with changes in walking behavior. Grip strength has often been associated with overall function and brain health in older adults, with greater grip strength correlating to greater overall strength, falls, and mortality (Bohannon, 2019; Carson, 2018; Pratama & Setiati, 2018). Pain perception and pain level are also associated with gait in older adults with chronic pain (Hicks et al., 2017; Kitayuguchi et al., 2016; Sawa et al., 2017; Taylor et al., 2018). Chronic pain has been associated with slower gait speed while walking on flat surfaces in older adults and while dual-tasking (Ogawa et al., 2020). Older adults with chronic pain may also develop fear avoidance and avoid physical activity to reduce or avoid pain (Camacho-Soto et al., 2012). This in part can also explain the previously found associations between slower gait speed and pain. Yet, we found that our measures of pressure pain threshold and grip strength did not significantly predict changes in walking speed during either dual-task or uneven terrain walking in our cohort of older adults. Future studies should consider adding other measures of pain (e.g., self-reported questionnaires) and sensorimotor function such as maximum voluntary contraction of lower extremities (Schantz et al., 1989). If these additional measures show differences across mobility function groups, it could further highlight why LFOA have decreased mobility across uneven terrains.

Alongside our sensorimotor predictors, the cognitive regression models also showed that changes in walking speed caused by uneven terrains were mainly predicted by NIH Flanker task performance, which reflects executive function including attention and inhibition. This association between NIH Flanker task and change in uneven terrain walking speed remained even when we accounted for mobility function with SPPB scores and sex differences, further inferring that changes to walking speed caused by uneven terrains were not only resultant of mobility function and sensorimotor function, but also by cognitive function as well. A recent study of obstacle walking by Chatterjee et al. (2020) also showed that older adults with lower executive function (quantified with a Trail Making Task and NIH EXAMINER executive function score) had greater decreases in obstacle walking speed compared to older adults with higher executive function. Furthermore, older adults with greater decreases in obstacle walking speeds also had lower prefrontal brain activity during walking, suggesting decreased capacity for cognitive compensation for increased walking difficulty. Within our current results, our finding suggests that walking on uneven terrains is modulated by attention and inhibitory function in older adults. Attention modulates processing of information from the walking environment (Hawkins, 1991; O’Connor et al., 2002) and influences walking behavior to adjustments in response to challenges in the environment, such as those faced with uneven terrains in the walking path. There is also accumulating evidence that poorer cognitive function, such as attention and working memory, is associated with higher fall risk (Montero-Odasso et al., 2012; Springer et al., 2006; Zhang et al., 2020). The cognitive and sensorimotor challenges posed by uneven terrain navigation may inherently increase this risk as older adults may have insufficient neural resources to control sensorimotor coordination.

As inferred by the results of the regression models and the changes in walking speed, our parametric increase in walking difficulty may impose cognitive challenges in addition to sensorimotor challenges, decreasing walking stability and therefore leading to slower walking speeds. A framework that may be helpful for interpreting the present results for parametric manipulation of task difficulty is the compensation related utilization of neural resources hypothesis (CRUNCH). CRUNCH is a neurocognitive model that addresses age differences in brain activity in response to varying levels of task demand, and associated effects on performance. The model conceptualizes how older adults exhibit differential task-related brain activity than young adults while performing tasks of increasing difficulty (Cappell et al., 2010; Reuter-Lorenz et al., 1999; Reuter-Lorenz & Cappell, 2008). In a working memory task, Cappell et al. (2010) showed that at lower levels of a working memory demand, older adults showed greater brain activity (e.g., bilateral prefrontal activations) compared to younger adults while maintaining comparable levels of task performance. However, at higher levels of task difficulty, this brain activity plateaued or decreased compared to young adults. This reduced available range of brain activity was associated with reduced task performance in older adults. According to the CRUNCH model, greater brain activation in older adults indicates some ability to compensate for age-related declines in brain structure and connectivity at low levels of task demand, but this ability reaches a ceiling at high levels of task difficulty when compensation is no longer possible. CRUNCH has not been investigated thoroughly in walking tasks where task difficulty is parametrically increased, such as walking on uneven terrains. Initial investigations in a visuomotor grasp task by Gerver et al. (2019) showed that, in older adults, performance of the grasp task declined as task difficulty was increased. Along with the decrease in motor performance, they observed increases in brain activity across frontoparietal regions. The same study reported a similar pattern of performance declines and increased brain activity in frontal brain regions when participants performed a working memory task. Van Ruitenbeek et al. (2022) recently showed that in a bimanual cursor tracking task, parametrically increasing the cursor tracking difficulty differentially affected brain activity between younger and older adults. As task difficulty increased, older adults showed increased brain activity at earlier task levels. However, older adults displayed a capability to increase their brain activation even with the highest task difficulties, contrary to the CRUNCH model. For their particular motor task, older adults did not reach a CRUNCH type ceiling. It is possible that the difficulty of the cursor tracking task did not reach a demand level where older adults failed to compensate for age-related declines. In recent work by our team (Downey et al., 2022), young adults and a subset of the older adults from this study performed uneven terrain walking on a treadmill, which resulted in greater perceived walking instability and increased gait variability in older adults as terrain unevenness was parametrically increased. Our team plans to extend the work of Downey and colleagues in future analyses of data collected within the Mind in Motion study (Clark et al., 2020). We aim to highlight the neural correlates of uneven terrain walking through mobile EEG recordings of brain activity and determine whether these correlates show CRUNCH related patterns of brain activity during increasing terrain unevenness. The results of these investigations will further highlight age-related changes to cognition and potential cognitive and/or sensorimotor targets for training to maintain walking performance in older adults.

The ultimate goal of our work to is elucidate new methods to best capture changes in walking behavior in older adults. Dual-task walking has provided unique insights into how varying attentional demands are met by individuals depending on cognitive status (e.g., executive resources/ability) and mobility level. The clinical applicability of dual-task walking tests has been well-explored in existing literature. For example, Yang and colleagues (2020) have published a comprehensive article on the reliability and validity of dual-task walking tests in people with chronic stroke. In another validity study, Muhaidat and colleagues (2014) showed that dual-task tests may be a useful tool to predict falls in older adults. Dual-task test results can also be used in clinical settings to determine gait stability and potential benefits from gait therapy (Lamoth et al., 2011; Tajali et al., 2019). In contrast, literature exploring the clinical relevance of walking in uneven terrain is scarce. Uneven terrain walking may also provide an alternative measurement of walking capability in daily life than flat, overground or treadmill walking typically done in the laboratory. That is, uneven terrain walking in the lab may be more ecologically relevant to study in older adults due to the rate of fall incidents around uneven terrain (Ambrose et al., 2013; Berg et al., 1997). Dual-task walking may also serve as an index of real-world walking providing that the concurrent task is calibrated sufficiently to match real-world scenarios such as walking on uneven terrains while talking or using a cellphone. This type of combined complex walking should be studied in the future to quantify age-related changes to walking behavior while walking in an environment that mimics everyday life. Existing studies have examined and confirmed the validity and clinical applicability of tools and scales that include a series of questions about walking on uneven terrains, such as the Lower Extremity Function Scale (Binkley et al., 1999) or the Cumberland Ankle Instability Tool (Hiller et al., 2006), but validity and reliability studies for clinic-conducted walking tests on uneven terrains are lacking. Even though more studies are necessary to determine the usability of our findings in a clinical setting, they can be used to equip clinicians and health practitioners with knowledge about the challenges older adults with lower mobility face when walking on uneven terrains and can have implications in fall prevention protocols. Berg et al. (1997) found that nearly 60% of falls are caused by trips and slips near uneven surfaces. A failure to adapt quickly to uneven terrains in the walking path could result in an increased rate of falls around these terrains. Future studies should consider a longitudinal follow-up to determine how uneven terrain walking behavior and adaptability is associated with future fall incidents for older adults.

### 4.1 Limitations

A limitation to the current study is that we did not obtain baseline measures of verbal fluency (in the absence of walking) which could characterize individual differences in this ability and assess tradeoffs when accompanied by walking. Nor did we manipulate task difficulty for the dual task walking condition, which prevents us from examining a slope of change in the same manner that we applied to uneven terrain walking. It is also possible that the level of demand imposed by the secondary cognitive task was already greater in magnitude as an intervention on gait dynamics compared to the three levels of uneven terrain magnitude. Thus, the comparison of walking speed changes across tasks could be limited by the differences in the magnitude of task difficulty. Although our manipulation across task types were not evenly distributed, we still determined how walking behavior changes as walking difficulty is increased across two walking tasks that differentially challenge walking, and identified predictors of these effects, providing insight to the underlying mechanisms.

## 5 Conclusions

In this study, we showed that in older adults, uneven terrains imposed differential costs (decreases) to walking speed from flat overground walking, compared to costs imposed upon walking speed by verbal dual-task walking. Uneven terrains in the walking path significantly decreased walking speed compared to walking on flat terrains and this effect was present across older adults with high and low mobility function. Older adults with poorer mobility function exhibited larger decreases in walking speed as terrain unevenness increased, compared to those with better mobility. We also found that selective attention and inhibitory function predicted changes in walking speed, across both dual-task and uneven terrain walking. Better cognitive performance in the NIH Flanker task was associated with smaller reductions in walking speed as walking difficulty increased. Changes to walking speed across both tasks were also predicted by 2PT discrimination thresholds, where better tactile discriminability corresponded to less reductions in walking speed. Overall, these findings suggest that uneven terrain and dual task walking present differential mobility challenges to older adults; further, uneven terrain is especially demanding for those with lower mobility function, poorer executive function, and worse tactile sensation.

## ACKNOWLEDGEMENTS

We thank all the participants for their time in the study and Megan Robertson, Nevena Stanojevic, and Lydia Mansour for participant recruitment, data collection, and data management.

## 1. Supplementary Materials

**Supplementary Table 1:**
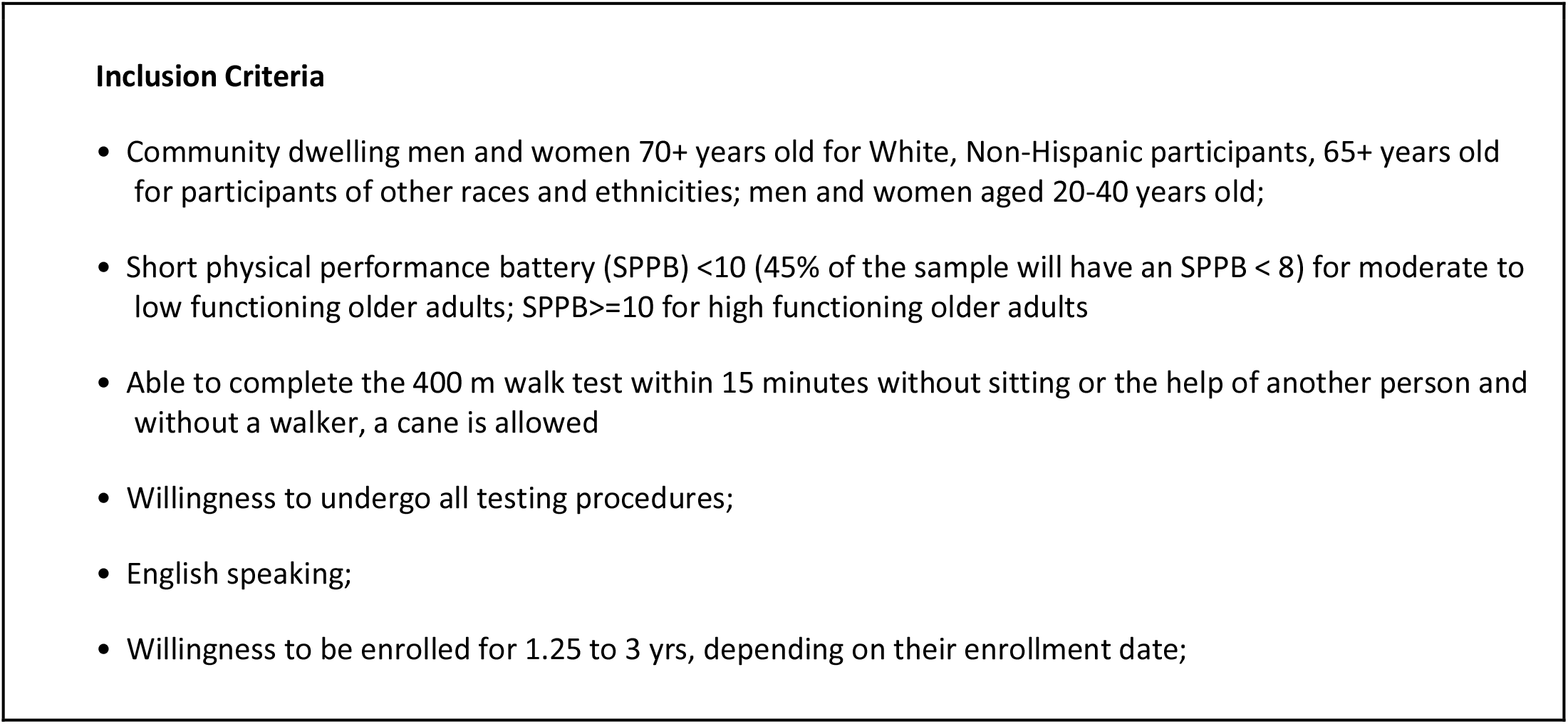
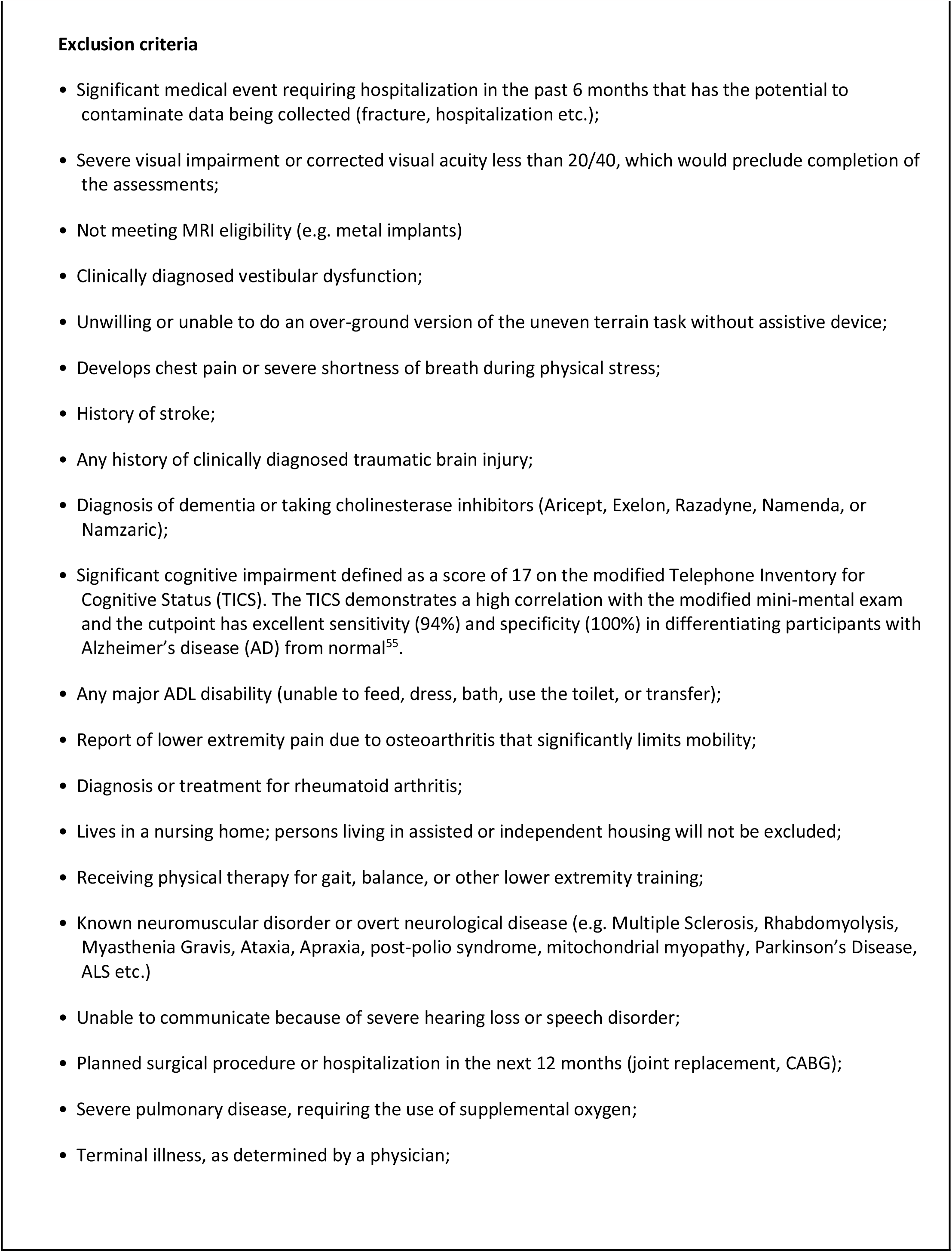
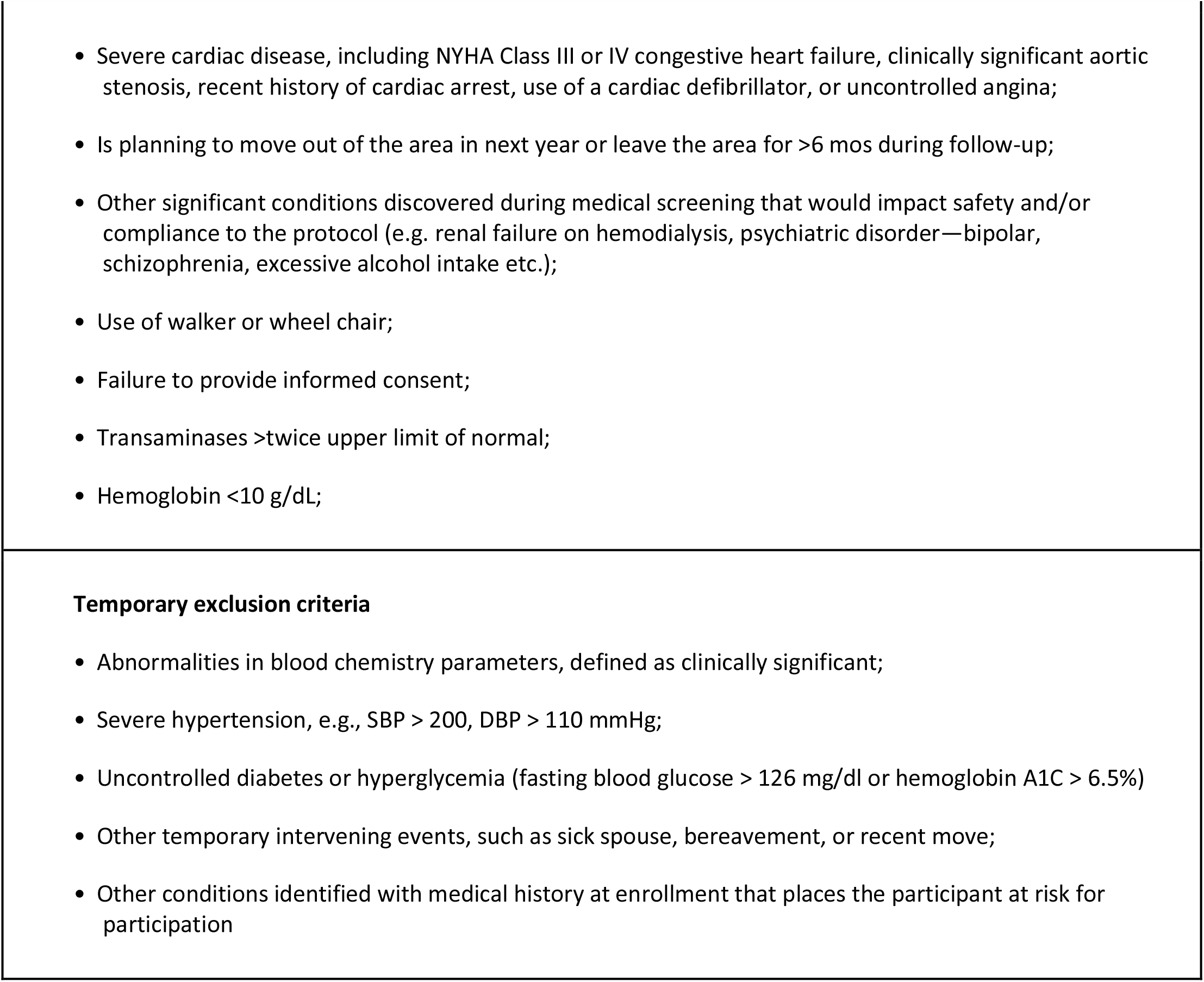
Inclusion and Exclusion Criteria for older adult cohort in the Mind in Motion study

**Supplemental Table 2:**
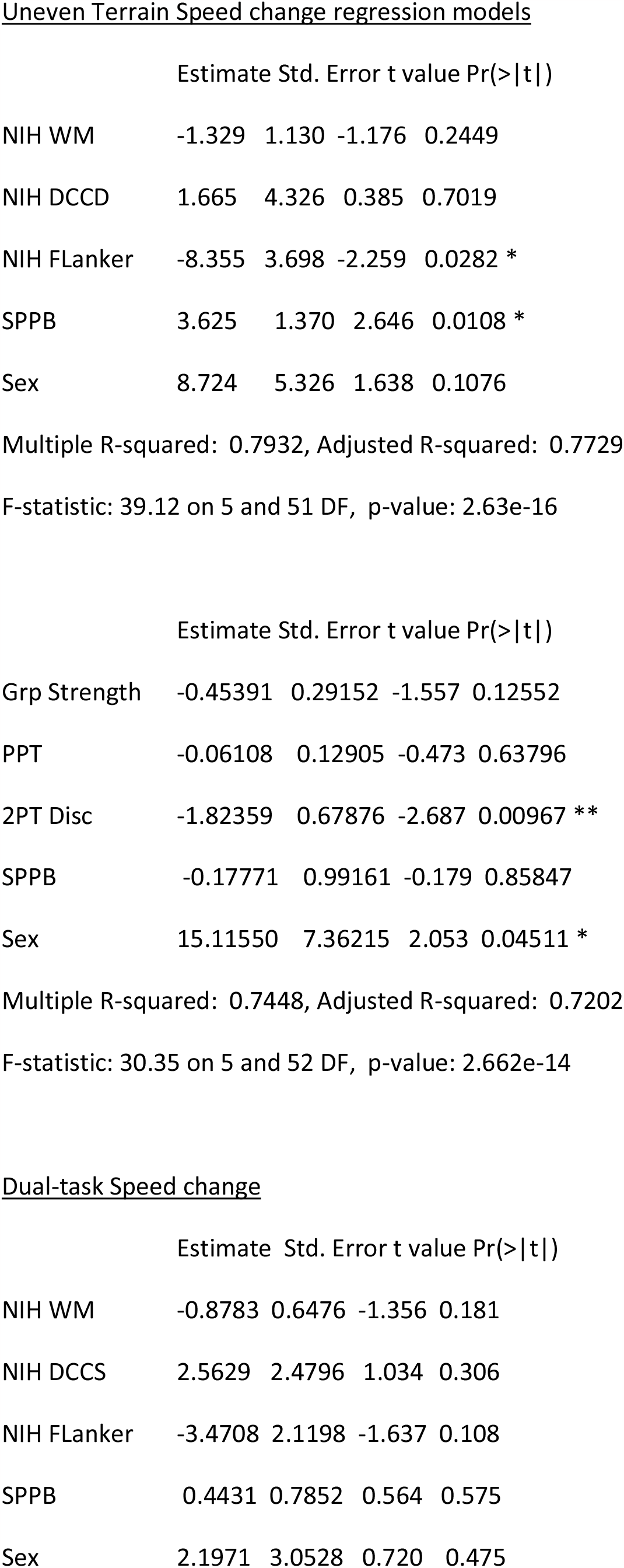
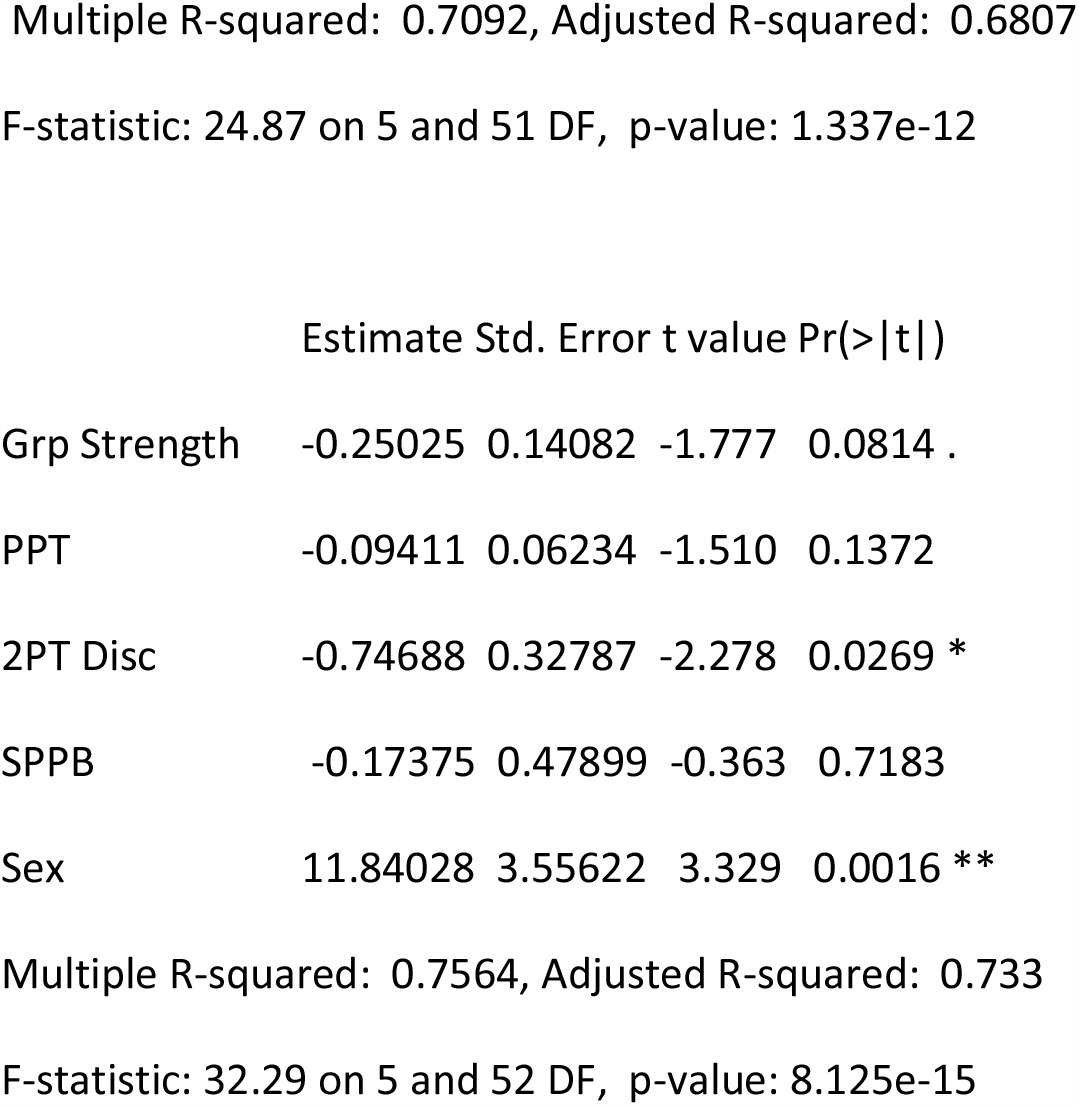
Regression analysis tables : Signif. codes: ‘***’ = p<0.001; ‘**’ = p<0.01; ‘*’ = p<0.05; ‘.’ = p<0.1

